# RGS6 regulates Kappa opioid receptor-mediated antinociceptive behaviors

**DOI:** 10.64898/2026.03.04.709600

**Authors:** Alyson P. Blount, Laurie P. Sutton

**Affiliations:** Department of Biological Sciences, University of Maryland Baltimore County, Baltimore, MD 21250, USA

**Keywords:** GPCR, Kappa opioid receptor (KOR), Anti-nociception, Regulators of G protein (RGS), Peripheral agonists, U50, 488, Analgesia

## Abstract

Targeting the kappa opioid receptor (KOR) system has emerged as a potential alternative to current analgesics, however, advancing the therapeutic development of KOR requires further elucidation of its intracellular signaling events and modulators. Among these intracellular modulators, Regulators of G protein signaling (RGS) proteins act as key modulators of GPCR signaling to shape nociceptive circuits and influence pain processing. Despite this, the molecular diversity of RGS proteins that shape KOR signaling and its behavioral consequences remains largely unexplored. Here we report that RGS6, a member of the R7 RGS family, is highly expressed in nociceptive areas and modulates multiple modalities of KOR-dependent anti-nociception and nocifensive behaviors. Using global single and double knockout mouse models we show that this anti-nociceptive phenotype was highly specific to RGS6 within the R7 RGS family. Further we demonstrate that the R7 RGS family displays a lack of functional redundancy in regulation of KOR signaling and behaviors. Using peripherally restricted KOR agonists, we found that KOR-RGS6 anti-nociceptive signaling displays sex differences in a site-specific manner, as females but not males displayed enhanced anti-nociceptive and blunted nocifensive behaviors. Our findings suggest that RGS6 is a highly specific modulator of KOR-dependent anti-nociceptive signaling and plays an essential role in modulating nociceptive circuits, potentially aiding in the development of novel analgesic drugs and therapeutics.

## INTRODUCTION

Pain is a multidimensional and universal process that serves as a critical protective mechanism that enables organisms to detect and respond to potentially harmful stimuli. However, dysregulation of pain signaling can lead to maladaptive pain states, where it has high comorbidity with mood disorders, substance abuse, and suicide (Best et al., 2022). Opioids are the gold standard therapeutic for the treatment of acute and chronic pain, with around 125 million opioids prescriptions distributed in the United States in 2023 alone (Khan et al., 2025). Long term use of opioids, however, can lead to serious and addictive side effects including respiratory depression, dependence, and overdose. Most clinically used opioid analgesics are selective agonists at the μ-opioid receptor (MOR) but the kappa opioid receptor (KOR) also modulates nociceptive transmission. In fact, the KOR and its endogenous ligand, dynorphin has increasingly become a target of interest as a potential alternative to current analgesics as it is nonaddictive, and lacks the respiratory failure typically associated with current opioid analgesics (El Daibani et al., 2024; Zhou et al., 2013).

KOR is a G protein coupled receptor (GPCR) that signals via both G protein and a β-arrestin pathways. Activation of KOR recruits inhibitory heterotrimeric G proteins which dissociates into Gα_i/o_ and Gβγ subunits upon GDP-GTP exchange at the Gα subunit. β-arrestin can also interact with KOR to promote the activation of several kinases. Early work has demonstrated that activation of G protein signaling by KOR elicits anti-pruritic and anti-nociceptive effects (Crowley and Kash, 2015; Liu et al., 2019; White et al., 2015). Subsequent β-arrestin signaling has been linked to KOR-induced aversion and sedation (Dunn et al., 2019; Ehrich et al., 2015). In recent years, there has been a growing interest in developing KOR selective G protein biased drugs for its analgesic properties. This in turn, has highlighted the need for a more detailed understanding of the G protein signaling events that mediate KOR-dependent anti-nociceptive properties.

Termination of G protein signaling is dependent on the inherent GTPase activity of the Gα_i/o_ protein and Regulators of G Protein Signaling (RGS) proteins enhance this GTPase activity. RGS proteins are GTPase activating proteins (GAPs) that quicken the hydrolysis rate of GTP on the Gα subunit, allowing the reformation of the inactive heterotrimeric G protein state, therefore influencing the physiologically relevant timing of G protein signaling (McPherson et al., 2018). RGS proteins ultimately cause inhibition of G protein signaling, decreasing its duration and thus are important regulators of G protein signaling cascades. Several RGS proteins have previously been implicated to modulate opioid anti-nociceptive signaling (Gross et al., 2019; Lamberts et al., 2013; Papachatzaki et al., 2011), however little is known on the molecular diversity of RGS proteins that shape KOR signaling and its behavioral consequences. The R7 RGS family is a main regulator of Gα_i/o_ coupled receptors and has been implicated to modulate nociceptive and MOR signaling (Garzón et al., 2003; Sutton et al., 2016; Zhou et al., 2012), making it an attractive potential modulator of KOR signaling. Two members of the R7 RGS family, RGS6 and RGS7, are widely expressed in the central nervous system (CNS) and have been indirectly implicated in regulation of KOR-induced behaviors. R7 family binding protein (R7BP), a membrane anchoring protein for the R7 RGS family, has been shown to be a modulator of KOR-induced anti-pruritis via the KOR, as it was found that a knockout (KO) of R7BP diminishes itch and is dependent on KOR signaling (Pandey et al., 2017). Further, a KO study of Gβ5, an obligate binding partner of the R7 RGS family, has implicated the family in modulation of multiple modalities of nociception, with a conditional KO of Gβ5 in RGS7 cells diminishing mechanical pain (Pandey et al., 2023). While indirect evidence suggests that RGS6 and RGS7 may modulate KOR signaling, their direct contributions to KOR-induced behaviors have yet to be examined. Identifying the individual R7 RGS family member that regulates KOR may elucidate the R7 RGS protein family’s *in vivo* contributions to KOR-mediated anti-nociception and related behaviors and may inform the development of KOR focused therapeutics. The goal of this study was to identify if the R7 RGS family modulates multiple modalities of KOR-induced anti-nociception, and if so, which member modulates KOR signaling. Here we identify a role of RGS6 on modulating KOR-induced anti-nociception and nocifensive behaviors to multiple modalities of nociception. This KOR-RGS6 modulation selectively influences anti-nociception without impacting KOR-induced sedation and is highly specific to RGS6 with no impact of RGS7 loss. Finally, the use of peripherally restricted KOR agonists induced anti-nociception and nocifensive behaviors in female RGS6 KOs but not males, implicating sex dependent modulation of KOR-RGS6 in a site-specific manner. RGS6 modulation of KOR signaling cascades may provide novel insights into the development of KOR-specific analgesic therapeutics.

## METHODS

### Animals

All studies were conducted in accordance with the National Institutes of Health (NIH) guidelines and were approved by the University of Maryland Baltimore County (UMBC) Institutional Animal Care and Use Committee regulations (IACUC). All animals were grouped housed under a 12/12 hour light-dark cycle with food and water available *ad libitum*. RGS7 knockout mice were generated as previously described and were kindly provided by Dr. Kirill A. Martemyanov (University of Miami) (Cao et al., 2012). RGS6 knockout mice were produced as previously reported and were generously provided by Dr. Kevin Wickman (University of Minnesota) (Yang et al., 2010). Breeding consisted of heterozygous matings, and all mice used were wildtype (^+/+^) and knockout (^-/-^) littermates. A double knockout (dKO) of RGS6 and RGS7 was generated by crossing homozygous RGS6^-/-^ to RGS7^-/-^ for two generations, with both dKOs and wildtype control littermates (RGS7^-/-^) obtained from RGS6 heterozygous RGS7^-/-^ breeding pairs. Male and female mice were used in the behavioral assays ranging from the age of 2-6 months.

### Drugs

KOR agonists (±)-U50,488 hydrochloride and ICI 404,488 hydrochloride were obtained from Tocris Bioscience (Bristol, UK). FE 200665 (i.e. CR665) was obtained from MedChem Express (Monmouth Junction, NJ, USA). All drugs were dissolved in 0.9% Saline and were administered intraperitoneally (IP) at doses specified.

### Behavioral Paradigms

Investigators were blinded to genotype and treatment during testing for all behavioral experiments conducted. All mice were habituated to the testing room for one hour before the start of any behavioral experiment.

#### Hot Plate

The hot plate (Ugo Basile, IT) was performed on a heated platform at 55^oC^ with a maximum cutoff time of 40 seconds. A pain response was recorded as the latency to lick/shake hind paws or jump. Once a pain response was observed or cutoff time was reached, mice were immediately returned to the homecage. The baseline pain response was recorded before drug treatment. An IP injection of U50,488 at 15 mg/kg was used to induce analgesia and the latency to a pain response was tested at 10, 30, 60, and 120 minutes post-injection. The peak analgesic response was calculated by subtracting the latency at 30 minutes by the baseline latency.

#### Cold Plate

The cold plate assay to measure cold-induced hyperalgesia was adapted from Madasu *et al*. (2021) using a Hot/Cold Plate from Ugo Basile, IT. Mice were treated with 5 mg/kg U50,488, 10 mg/kg ICI 204,488, 12 mg/kg FE 200665, or 0.9% saline and placed back into the homecage. 30 minutes following drug treatment, the mice were placed on the cold plate for 5 minutes, and the latency to jump and the number of jumps was recorded as nocifensive responses. The temperature of the plate was either 30^oC^, 10 ^oC^, or 3^oC^ based on the experiment.

#### Electronic Von Frey

To evaluate mechanical allodynia/hyperalgesia an electronic Von Frey (Ugo Basile) paradigm was utilized. Mice were habituated to the clear acrylic apparatus with a raised metal mesh floor (96mm x 96mm x 140mm; Ugo Basile, IT). A Von Frey metal probe was applied with increasing pressure to the midplantar surface of the hind paw. The minimum applied pressure (gf) at time of the paw retracting from the Von Frey probe was recorded. Baseline nociception was collected 24-48 hours before drug treatment. To obtain the Mechanical Withdrawal Threshold (MWT), which is the minimum amount of force that elicits a withdrawal reflex, 5 measurements were obtained with at least 5 minutes between measurements. The 3 values that deviated the least from the median were averaged with measurements below 4 gf removed. This was to ensure that all mice had a consistent and sufficiently high baseline. Mice were then treated with 15 mg/kg U50,488, 10 mg/kg ICI 204,488, or 12 mg/kg FE 200665 and measurements were taken every 60 minutes to 300 minutes. The MWT for each time point was measured 3 times and averaged for each mouse with 5 minutes allotted between measurements.

#### Rotarod Sedation Assay

Assessed motor coordination and sedation using an accelerating rotarod (Med Associates Inc., VT, USA) as previously reported (White et al., 2015). The rotarod program increased from 3 to 30 RPM over 5 minutes. Each trial ended when the mouse fell off the rod or reached 5 minutes and the time and RPM were recorded. Mice were trained up to 3 trials in one day and returned to homecage between trials. On testing day, baseline time on the rod and speed was recorded before drug administration. Mice received an IP injection of 15 mg/kg U50,488 and were assessed at 10, 30, 40, and 60 minutes on the rotarod.

### *In situ* Hybridization (RNAscope)

Fresh-frozen RNAscope protocol was adapted from Advanced Cell Diagnostics (ACD) website (https://acdbio.com/rnascope-multiplex-fluorescent-v2-assay) using the RNAscope™ Multiplex Fluorescent Reagent Kit v2 (ACD, Cat. No. 323100). The Nucleus Accumbens (NAc), Ventral Tegmental Area (VTA), Dorsal Root Ganglions (DRGs), and the cervical Spinal Cord (SC) were freshly dissected out of mice, embedded in OCT (Fisher Scientific) and flash frozen in liquid nitrogen. Tissues were sectioned (Leica CM 3050 S) at 20 µm, collected on Superfrost Plus slides (Fisher Scientific), and stored at -80^oC^ until use. Slides were washed in prechilled 4% PFA for 15 minutes, washed with 1X PBS and pretreated per manufacturer’s instructions using RNAscope H2O2 and Proteases Reagents kit (ACD, Cat. No. 322381). Target probes for *oprk1* (Cat. No 316111-C2), *rgs6* (Cat. No 521211), and *rgs7* (Cat. No 519291-C3) were combined and hybridized for 2 hours at 40^oC^ using the HybEZ Hybridization System (ACD, Cat. No. 321711). Following amplification, probes were fluorescently labeled with Opal 520, Opal 570, and Opal 690 (Akoya Biosciences) diluted at 1:1500. Slides were counterstained with DAPI or NeuroTrace 435/355 (1:50 dilution; Thermo Fisher Scientific, Cat. No. N21479) and mounted with ProLong Gold Antifade Mountant (Thermo Fisher Scientific). Slides were imaged with a Zeiss LSM 900 confocal microscope at 20X, 40x, and 63X objectives. Quantifications of images were performed in ImageJ Fiji. Cells with clear fluorescent puncta were defined as expressing targeted RNA.

### Quantification and Statistical Analysis

Statistical analysis was performed using GraphPad Prism (Prism 10.6.1, GraphPad Software). Data was analyzed using 2-way repeated measure (RM) ANOVA, 2-way ANOVA, or an unpaired 2-tailed t-test. Post hoc tests used were Turkey’s multiple comparisons or Fisher’s LSD unless otherwise specified. Differences were considered significant if p< 0.05. All data are expressed as mean ± SEM.

## RESULTS

### *Oprk1+* cells in pain related areas exhibit strong co-expression of *RGS6* and *RGS7*

We assessed co-expression of RGS6 and RGS7 in pain related areas with robust KOR expression, including the spinal cord (SC), the dorsal root ganglions (DRG), the nucleus accumbens (NAc), and the ventral tegmental area (VTA) (Cai et al., 2016). The mRNA expression of *rgs6* and *rgs7* in *Oprk1* positive (*oprk1+*) cells in C57BlJ/6 mice was assessed in the NAc, VTA, SC, and DRGs using a 3-plex RNAscope and was imaged at 20X, 40X, and 63X. In the NAc, it was found that of the *oprk1+* cells, 75.61% of them expressed both *rgs7* and *rgs6*, 11.79% expressed only *rgs6*, 4.47% expressed *rgs7* alone, and 8.13% had neither RGS protein mRNA **(Figure 1A, 1B)**. In the VTA, co-expression of *oprk1*, *rgs6*, and *rgs7* consisted of 63.45% of *oprk1+* cells, 4.83% of *oprk1+* cells were *rgs6* only, 16.55% with *rgs7* only, and 15.17% with only *oprk1+* **(Figure 1C, 1D).** Together, our data indicates that a high percentage of *oprk1+* cells co-expressed both *rgs6* and *rgs7* in supraspinal nociceptive areas. In the SC, 53.49% of *oprk1+* cells expressed both *rgs6* and *rgs7*, 24.03% of *oprk1+* cells showed co-expression of *rgs6* alone, 8.53% expressed *rgs7* alone, and 13.95% expressed neither R7 RGS member **(Figure 1E, 1F)**. When assessing co-expression in the DRGs, we used neurotrace to assess cell boundaries in the DRGs due to irregular cell sizes. *Oprk1+* cells in the DRGs co-expressed *rgs6*, and *rgs7* together in 42.57% of cells, *rgs6* alone in 17.57%, *rgs7* only in 25.68%, and co-expression with neither RGS protein in 14.19% of cells **(Figure 1G, 1H)**. All areas tested showed a large percentage of overlap between *oprk1* and the two R7 RGS proteins, implicating both RGS6 and RGS7 as being possible modulators of KOR signaling.

**Figure 1.**
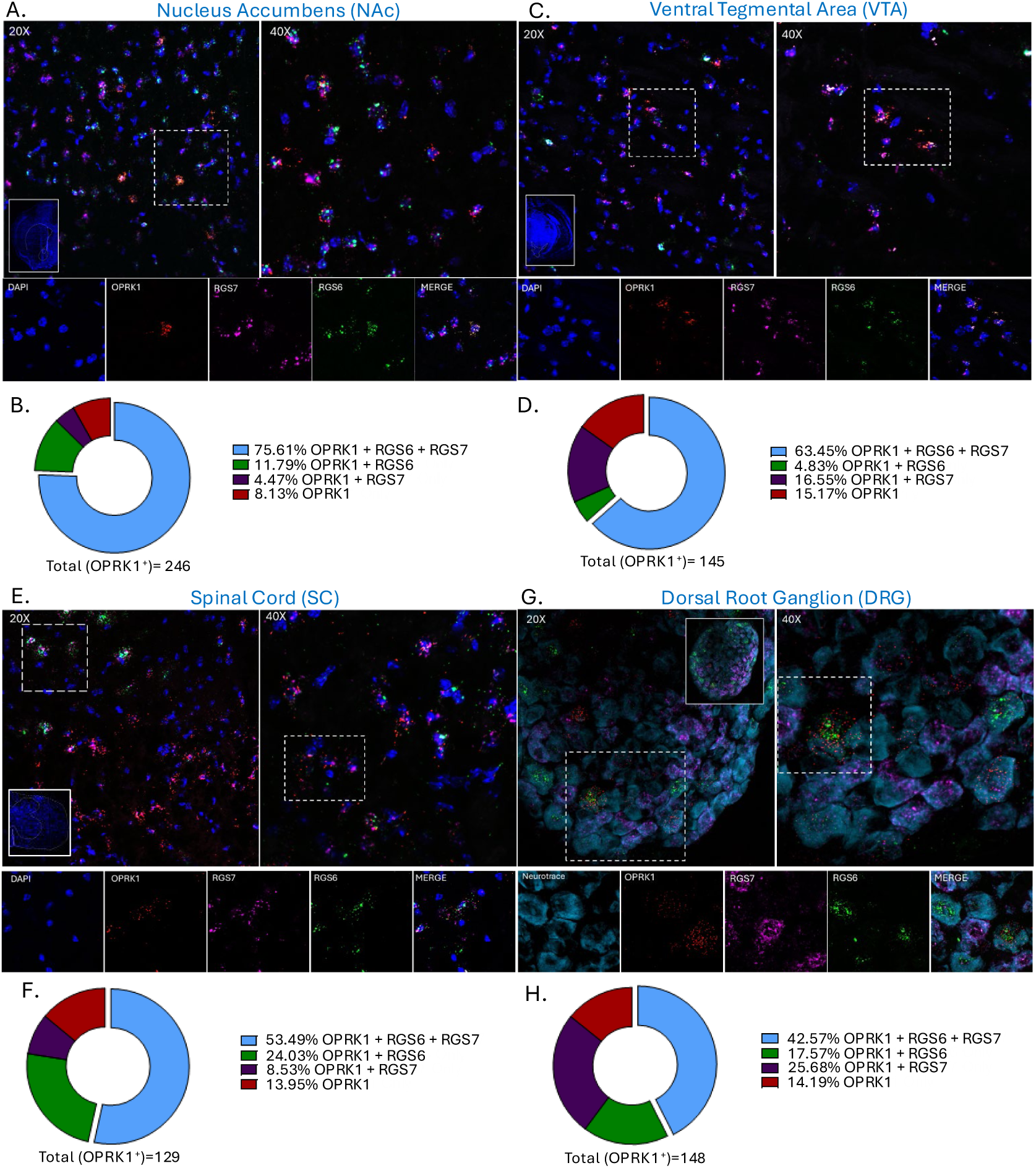
Expression of *rgs6* and *rgs7* mRNA in brain areas implicated in KOR-dependent behaviors. Representative RNAscope images of *oprk1*, *rgs6*, and *rgs7* at 20X, 40X, and 63X in the Nucleus Accumbens (NAc)(**A**), Ventral Tegmental Area (VTA) (**C**), Spinal Cord (SC) (**E**), and the Dorsal Root Ganglion (DRG) (**G**). Quantification of *oprk1*, *rgs6*, and *rgs7* mRNA transcript colocalization in the NAc (**B**), the VTA (**D**), the SC (**F**), and the DRGs (**H**). Inserts display representative images of examined areas (dashed lines).

### Lack of RGS6 enhances KOR-dependent anti-nociceptive responses to mechanical pain

To assess the contributions of RGS6 and RGS7 in KOR induced mechanical anti-nociception, we employed a von frey paradigm to assess mechanical withdrawal thresholds (MWT) using two transgenic knockout mouse models, RGS6^-/-^ and RGS7^-/-^ along with their wildtype RGS6^+/+^ and RGS7^+/+^ littermates **(Figure 2A)**. Previous work has shown sex differences in the effectiveness of KOR-induced thermal analgesia to multiple KOR agonists (Craft and Bernal, 2001; Stoffel et al., 2005), therefore, we assessed potential sex differences in KOR behaviors in both of our knockout models **(Figure 2B)**. We validated in both male and female C57BLJ6 mice that 10 mg/kg U50,488 was sufficient to induce a significant levels of anti-nociception as shown in an effect of time (F _(4, 164)_ = 57.71, p<0.0001), sex (F _(1, 41)_ = 7.242, p=0.0103), and time x sex (F _(4, 164)_ = 6.422, p<0.0001) **(Figure 2C)**. After injection, both male and female mice showed a peak in MWTs at 180 minutes, with female mice showing lower MWT compared to males at 180 minutes after injection, suggesting that the anti-nociceptive effect of U50,488 was more effective in males in a mechanical pain assay.

**Figure 2.**
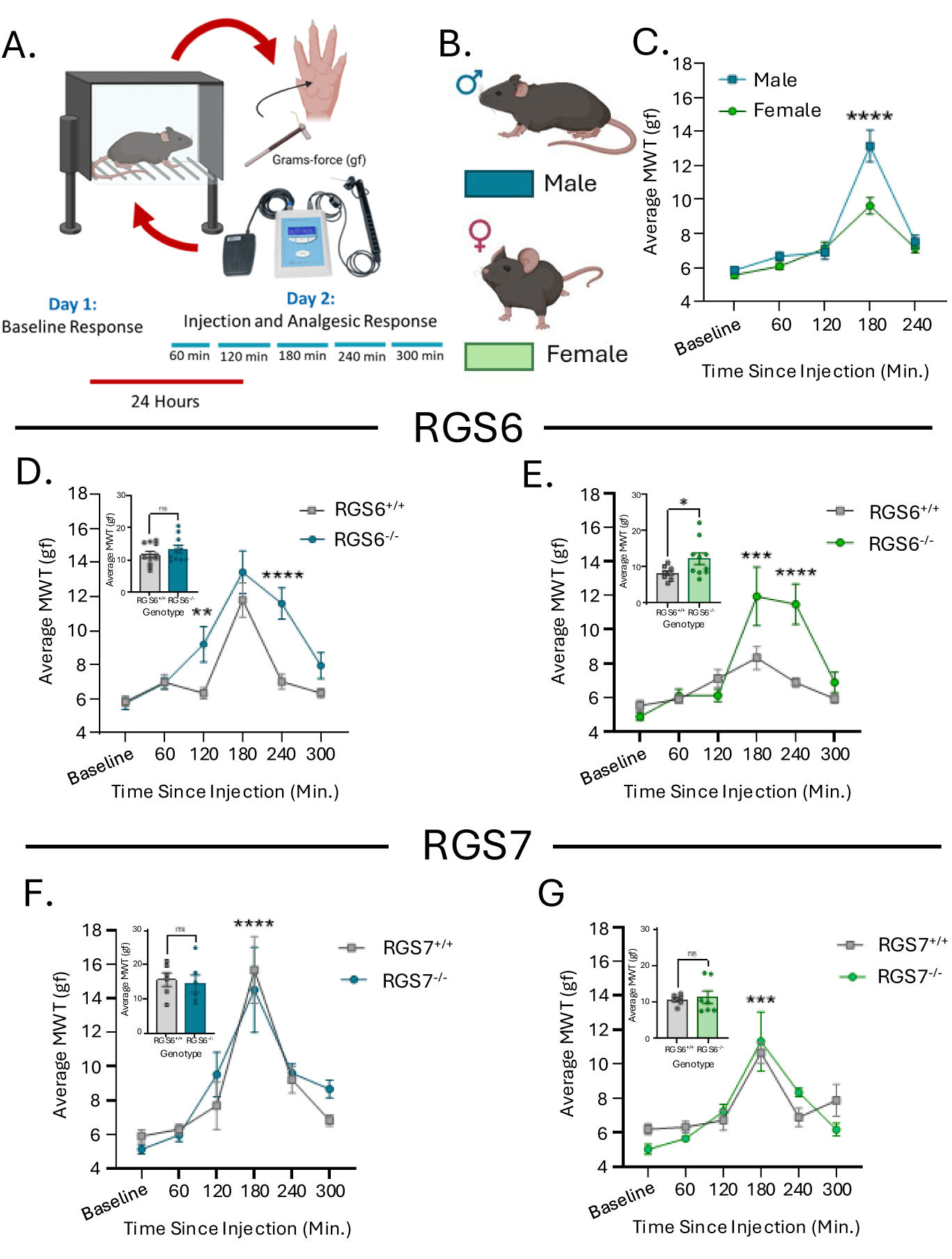
RGS protein family on mechanical pain using an Von Frey Assay and Kappa Opioid Receptor (KOR) unbiased agonist, U50,488. **A.** Timeline and schematic of the Von Frey procedure. **B.** Key for male and female mice. **C.** Mechanical Withdrawal Threshold (MWT) after an injection of over time in male (N=23) and female (N=20) C57BLJ6 mice. **D.** Average MWT after U50,488 injection over time in RGS6^+/+^ and RGS6^-/-^ male mice (N=11-10). Insert is the average MWT of male RGS6^+/+^ and RGS6^-/-^ at the peak analgesic response point of 180 minutes after injection. **E.** Average MWT after an injection U50,488 over time in RGS6^+/+^ and RGS6^-/-^ female mice (N= 8-10). Insert is the average MWT of female RGS6^+/+^ and RGS6^-/-^ 180 minutes after injection. **F.** Average MWT after an injection U50,488 over time in RGS7^+/+^ and RGS7^-/-^ male mice (N=6). Insert is the average MWT of male RGS7^+/+^ and RGS7^-/-^ at 180 minutes after injection. **G.** Average MWT necessary to elicit a pain response after an injection U50,488 over time in RGS7^+/+^ and RGS7^-/-^female mice (N=6-7). Insert is the average MWT of female RGS7^+/+^ and RGS7^-/-^ 180 minutes after injection. Data shown as means ± SEM; * p≤0.05, ** p≤0.01, *** p≤0.001, **** p<0.0001.

We assessed the MWT of male RGS6^+/+^ and RGS6^-/-^ mice overtime and found that lack of RGS6 induces significantly elevated MWT compared to RGS6^+/+^ mice (time F _(5, 95)_ = 24.97, p=<0.0001; genotype F _(1, 19)_ = 17.83, p=0.0005; interaction F _(5, 95)_ = 3.382, p=0.0074) **(Figure 2D).** Despite showing significantly higher levels of KOR-induced anti-nociception over time, male RGS6^-/-^ mice showed no difference from RGS6^+/+^ at the peak analgesic time point of 180 minutes **(Figure 2D insert)**. It is possible that there is a ceiling effect at 180 minutes for both RGS6^+/+^ and RGS6^-/-^ mice, as male RGS6^-/-^ mice show a phenotypic difference at time points on either side of the peak analgesic time. Female RGS6^-/-^ mice in this assay also showed enhanced anti-nociceptive effects to mechanical pain when treated with U50,488 compared to RGS6^+/+^ mice (time F _(5, 70)_ = 14.32, p<0.0001; genotype F _(1, 14)_ = 12.65, p=0.0005; interaction F _(5, 70)_ = 5.032, p<0.0001) **(Figure 2E).** Female RGS6^-/-^ mice showed heightened KOR-induced anti-nociception at 180 minutes (t _(16)_ =2.138, p=0.0482) **(Figure 2E inset**), which may be attributed to the decreased effectiveness of KOR anti-nociception in RGS6^+/+^ female mice.

RGS7^+/+^ and RGS7^-/-^ mice were similarly assessed. Male RGS7^+/+^ and RGS7^-/-^ mice had significantly increased MWT at comparable levels at 180 minutes after injection from baseline (time F _(5, 50)_ = 22.06, p<0.0001) with no effect of genotype over time (F _(1, 10)_ = 0.1082, p=0.7490; insert t _(10)_ =0.3670, p=0.7212) **(Figure 2F, insert)**. Female RGS7^+/+^ and RGS7^-/-^ mice treated with U50,488 both show increased MWT (time F _(5, 55)_ = 14.72, p<0.0001) with no effect of genotype at the peak analgesic time point (F _(1, 11)_ = 0.1621, p=0.6949; insert t _(11)_ =0.3302, p=0.7475) **(Figure 2G, insert).** Our data suggests that RGS6 but not RGS7 may modulate KOR-induced anti-nociception in response to mechanical pain.

### Knockout of RGS6 enhances KOR-dependent thermal anti-nociception

Due to enhanced KOR-induced mechanical anti-nociception displayed in our RGS6^-/-^model, we investigated the effect of a lack of RGS6 and RGS7 on thermal KOR anti-nociception **(Figure 3A)**. To validate the established sex difference present in KOR-induced analgesia, we recapitulated previous results implicating differential effects of sex on KOR-dependent thermal analgesia (time F _(4, 140)_ = 7.947, p<0.0001; sex (F _(1, 35)_ = 9.769, p=0.0036); interaction (F _(4, 140_ = 6.766, p=0.0001) **(Figure 3B)**. Comparison of male RGS6^+/+^ and RGS6^-/-^ mice revealed that while both genotypes show an increase in latency to show a pain response on the hot plate (time F _(4,68)_ = 21.88, p<0.0001), RGS6^-/-^ mice display a significantly higher peak analgesic response compared to RGS6^+/+^ mice 30 minutes after U50,488 injection (genotype F _(1, 17)_ = 4.832, p=0.0421) **(Figure 3C).** Female RGS6^-/-^ mice also show an increased latency compared to RGS6^+/+^ mice at the peak response time (genotype F _(1, 14)_ = 4.847, p=0.0450) with an effect of time (F _(4, 56)_ = 2.564, p=0.048) **(Figure 3D).** The peak analgesic response of KOR-induced anti-nociception between male and female RGS6^+/+^ and RGS6^-/-^ mice was compared and displayed an effect of sex (F _(1, 30)_ = 19.10, p=0.0001) and genotype (F _(1, 30)_ = 11.26, p=0.0022) **(Figure 3E)**. The peak thermal analgesic response reveals that both sexes of RGS6^-/-^ mice display an increased anti-nociceptive response compared to RGS6^+/+^ mice, but the sex difference in KOR-induced anti-nociception was still maintained, with the female RGS6^+/+^ mice showing less of an effect compared to males.

**Figure 3.**
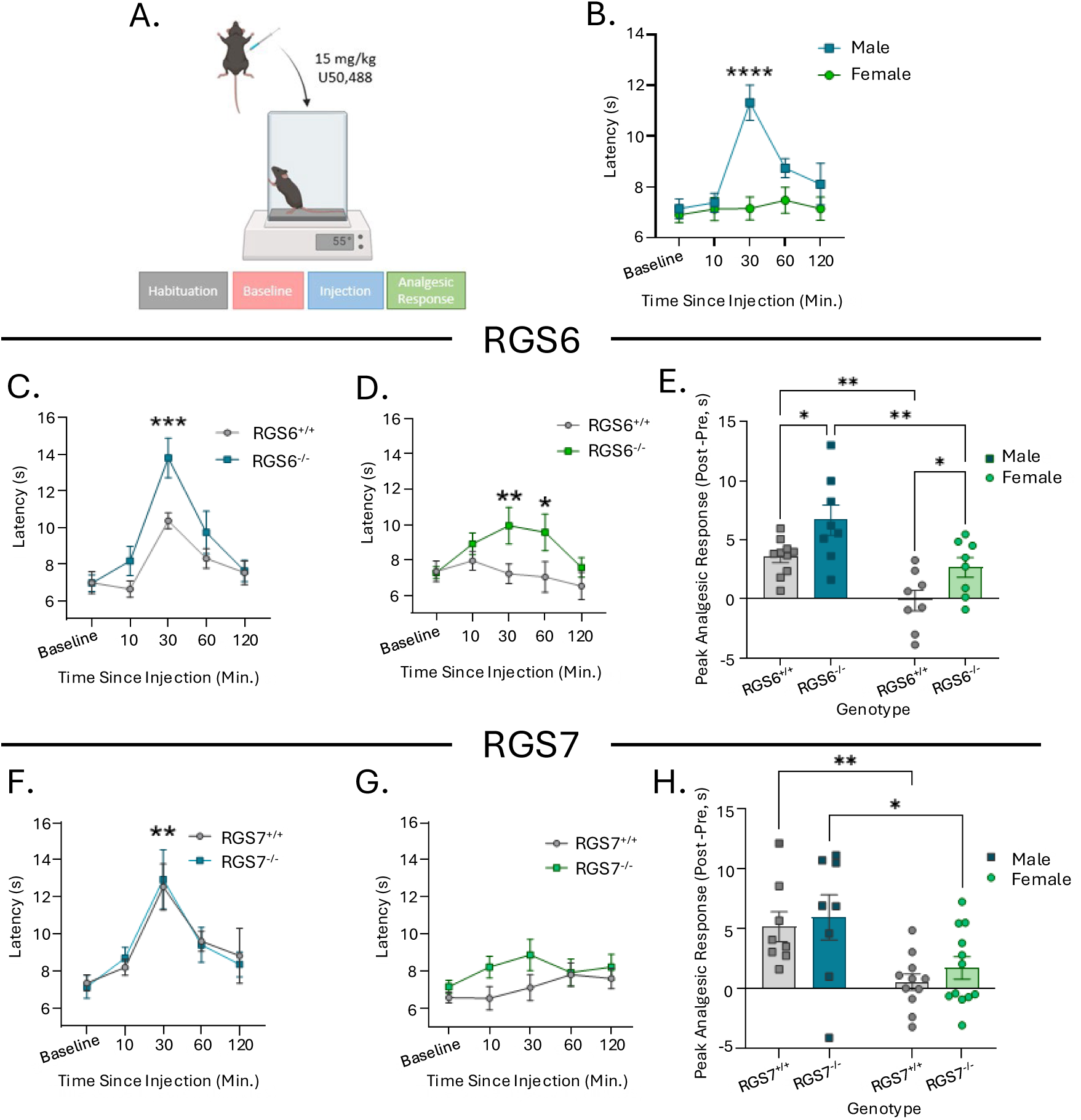
Role of R7 RGS protein family on KOR-induced thermal analgesia. Colors assigned to males and females are blue and green respectively. **A.** Schematic and timeline of hot plate with U50,488 (IP) in both sexes. **B.** 15 mg/kg U50,488 on KOR-induced analgesia in C57BLJ6 male and female mice (N=18-19). **C.** Average latency to show a pain response over time in male RGS6^+/+^ and RGS6^-/-^ mice (N=10-9). **D.** Average latency to show a pain response over time in female RGS6^+/+^ and RGS6^-/-^ mice (N=8). **E.** Comparison of peak analgesic response between RGS6^+/+^ and RGS6^-/-^ in male and female mice. **F.** Average latency to show a pain response over time in male RGS7^+/+^and RGS7^-/-^ mice (N=8). **G.** Average latency to show a pain response over time in female RGS7^+/+^and RGS7^-/-^ mice (N=11-12). **H.** Comparison of peak analgesic response between RGS7^+/+^and RGS7^-/-^ in male and female mice. Data shown as means ± SEM; * p≤0.05, ** p≤0.01, **** p≤0.001.

Both male RGS7^+/+^ and RGS7^-/-^ mice demonstrated KOR-dependent analgesic effects with no significant effect on genotype (time F _(4, 64)_ = 10.78, p<0.0001; genotype F _(1, 16)_ = 0.0002749, p=0.987) **(Figure 3F)**. No analgesic effect was induced in either genotype of RGS7 female mice (genotype F _(1, 21)_ = 2.814, p=0.1083, time F _(4, 84)_ = 1.578, p=.0.1876) **(Figure 3G)**. Comparison of the peak analgesic response between both sexes of RGS7^+/+^ and RGS7^-/-^ mice, showed a significant effect of sex (F _(1, 35)_ = 14.26, p=0.0006) but not genotype (F _(1, 35)_ = 0.7004, p=0.4083), demonstrating that RGS7 does not modulate KOR-dependent thermal anti-nociception **(Figure 3H)**.

### Lack of RGS6 elicits blunted KOR-dependent noxious cold hypersensitivity and nocifensive responses

KOR activation is involved in driving noxious cold hypersensitivity and inducing nocifensive behavior, however it is unknown if the R7 RGS family mediates this effect. To assess if the R7 RGS family may modulate this effect, we tested KOR-induced cold hypersensitivity and nocifensive responses in RGS6^-/-^ and RGS7^-/-^ mouse models. Excessive jumping behavior is commonly classified as a nocifensive response to noxious stimuli, including noxious cold (Deuis et al., 2017). To validate this model, we measured the latency to jump and the number of jumps on the cold plate for 5 minutes in male and female C57BL6J mice across a temperature gradient **(Figure 4A)**. Male C57BL6J mice that received an injection of U50,488 showed a significantly higher amount of jumps in response to 3^oC^ compared to baseline (temperature F _(2, 36)_ = 9.001, p=0.0007; treatment F _(1, 36)_ = 19.78, p<0.0001; interaction F _(2, 36)_ = 7.170, p=0.0024) **(Figure 4B)**. A similar effect was seen in female C57BL6J mice (temperature F _(2, 40)_ = 15.91, p<0.0001; treatment F _(1, 40)_ = 22.26, p<0.0001; interaction F _(2, 40)_ = 10.63, p=0.0024) **(Figure 4C)**. At 30^oC^ or 10^oC^ neither sex displayed differences in the number of jumps between baseline and U50,488 treated mice. This was validated when assessing latency, as there was a significant decrease in the latency to jump in both male (temperature F _(2, 37)_ = 7.821, p=0.0015; genotype F _(1, 37)_ = 13.35, p=0.0008) **(S. Figure 1A)** and female C57BLJ6 mice (temperature F _(2, 41)_ = 7.253, p=0.002; genotype F _(1, 41)_ = 7.544, p=0.009) **(S. Figure 1B**) upon U50,488 treatment at 3^oC^. We found that while hypersensitivity was induced in both sexes of C57BL6J, females show a mild increased sensitivity to KOR induced hypersensitivity at noxious temperatures (t _(14)_ =2.348, p=0.0341) **(Figure 4D).**

**Figure 4.**
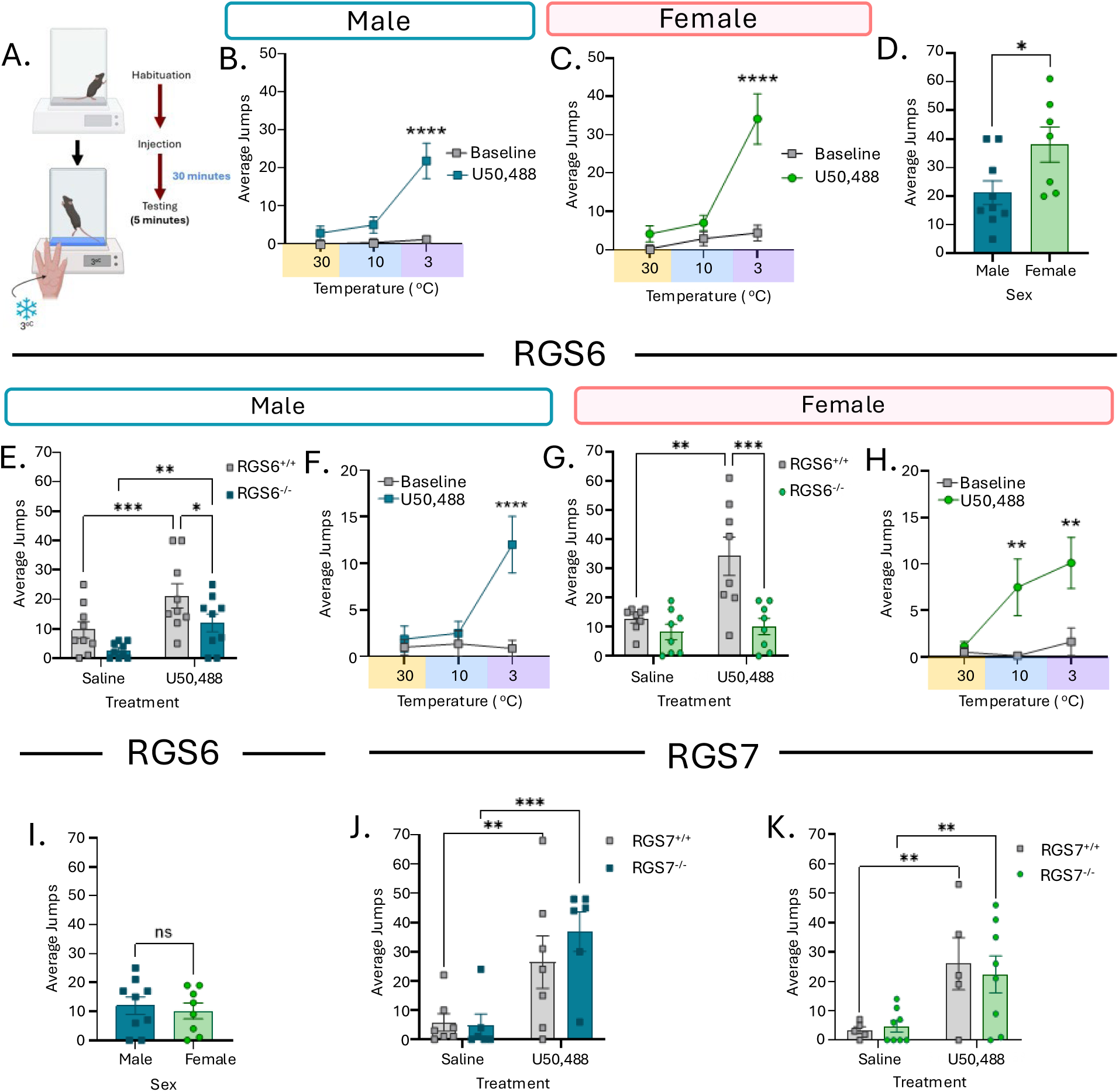
RGS6 and RGS7 on KOR-induced noxious cold hypersensitivity. **A.** Depiction of the cold plate experimental procedure. Mice were injected with either 0.9% Saline or a KOR agonist. **B.** Average number of jumps in male C57BLJ6 mice when treated with 5 mg/kg U50,488 compared to baseline on a temperature gradient of 30^oC^, 10^oC^, & 3^oC^ (N=9). **C.** Average number of jumps in female C57BLJ6 mice when treated with U50,488 compared to baseline on a temperature gradient of 30^oC^, 10^oC^, & 3^oC^ (N=7). **D.** Comparison of the average number of jumps at 3^oC^ between C57BLJ6 male and female mice. **E.** Average number of jumps on a 3^oC^ cold plate after an injection of saline or U50,488 in male RGS6^+/+^ and RGS6^-/-^ mice (N=9). **F.** Average number of jumps in male RGS6^-/-^ mice when treated with U50,488 compared to baseline on a temperature gradient of 30^oC^, 10^oC^, & 3^oC^. **G.** Average number of jumps on a 3^oC^ cold plate after an injection of saline or U50,488 in female RGS6^+/+^ and RGS6^-/-^ mice (N=8). **H.** Average number of jumps in female RGS6^-/-^ mice when treated with U50,488 compared to baseline on a temperature gradient of 30^oC^, 10^oC^, & 3^oC^. **I.** Comparison of the average number of jumps at 3^oC^ between RGS6^-/-^ male and female mice (N=9-8). **J.** Average number of jumps on a 3^oC^ cold plate after an injection of saline or U50,488 in male RGS7^+/+^ and RGS7^-/-^ mice (N=7). **K.** Average number of jumps on a 3^oC^ cold plate after an injection of saline or U50,488 in female RGS7^+/+^ and RGS7^-/-^ mice (N=5-9). Data shown as means ± SEM; * p≤0.05, ** p≤0.01, *** p≤0.001, **** p<0.0001.

To assess if RGS6^+/+^ and RGS6^-/-^ mice modulate KOR-induced cold hypersensitivity and nocifensive jumping behavior, mice treated with saline or U50,488 were exposed to 3^oc^ and nocifensive jumping was assessed. Male RGS6^+/+^ and RGS6^-/-^ mice showed a significant increase in the number of jumps indicating KOR hyperactivity in response to U50, 488 (treatment F _(1, 16)_ = 26.94, p<0.0001), however RGS6^-/-^ mice showed a blunted nocifensive jumping response compared to RGS6^+/+^ mice (genotype F _(1, 16)_ = 4.928, p=0.0412) **(Figure 4E).** When assessing the latency to jump, male RGS6^-/-^ mice show a trend towards increased latency compared to RGS6^+/+^, supporting blunting of KOR-induced cold nocifensive (t _(17)_ =1.621, p=0.1234) **(S. Figure 1C)**. To see if this effect was specific to 3^oC^, we evaluated male RGS6^-/-^ mice on a temperature gradient, where male U50,488-treated RGS6^-/-^ mice exhibited a significant increase in nocifensive behavior at 3^oc^ compared to 30^oc^ and 10^oc^ (temperature F _(2, 42)_ = 5.029, p=0.011; treatment F _(1, 42)_ = 9.266, p=0.004; interaction F _(2, 42)_ = 5.686, p=0.006) **(Figure 4F)**. When further examined on a temperature gradient, male U50,488-treated RGS6^-/-^ mice displayed decreased latency to jump compared to baseline jumping in RGS6^-/-^ mice at 3^oc^ (treatment F _(1, 44)_ = 4.765, p=0.0344; interaction F _(2, 44)_ = 4.953, p=0.0115) **(S. Figure 1D)**. In female RGS6^+/+^ and RGS6^-/-^ mice exposed to 3^oc^ cold plate, we found that U50,488 induces elevated nocifensive jumps from saline in RGS6^+/+^ but not RGS6^-/-^ mice, with RGS6^-/-^ mice revealing blunted KOR-induced cold hypersensitivity (treatment F _(1, 14)_ = 9.267, p=0.0088; genotype F _(1, 14)_ = 13.23, p=0.0027; interaction F _(1, 14)_ = 6.518, p=0.0230) **(Figure 4G).** This was supported by the change in latency to jump (t _(13)_ =4.012, p=0.0015) **(S. Figure 1E)**. When female RGS6^-/-^ mice were assessed on a temperature gradient, we found increased nocifensive jumping at 3^oc^ and 10^oc^ compared to saline (treatment F _(1, 41)_ = 13.31, p=0.0007; temperature F _(2, 41)_ = 3.723, p=0.0327) **(Figure 4H)**. A similar effect in latency was observed as U50,488 RGS6^-/-^ mice show a decreased latency to jump at 3^oc^ and 10^oc^ compared with saline treated (temperature F _(2, 41)_ = 4.178, p=0.02; treatment F _(1, 41)_ = 18.66, p<0.0001) **(S. Figure 1F)**. This demonstrates that the lack of RGS6 blunts the response to noxious cold in both sexes (t _(15)_ =0.4542, p=0.6562) **(Figure 4I).**

To determine if RGS7 plays a role in KOR-induced hypersensitivity to cold, we tested male RGS7^+/+^ and RGS7^-/-^ mice on a 3^oC^ cold plate assay. Male RGS7^+/+^ and RGS7^-/-^ mice treated with U50,488 both show significant KOR-dependent nocifensive responses compared to saline with no effect of genotype (treatment F _(1, 11)_ = 29.36, p=0.0002; genotype F _(1, 11)_ = 0.4033, p=0.5384) **(Figure 4J).** The latency of cold induced nocifensive jumping also showed no effect of genotype (t _(11)_ =0.2007, p=0.8446) **(S. Figure 1G).** Female RGS7^+/+^ and RGS7^-/-^ mice show similar effects, with an increase in jumps to U50,488 (treatment (F _(1, 11)_ = 19.39, p=0.0011) but no difference between genotypes (F _(1, 11)_ = 0.03338, p=0.8584) **(Figure 4K)**. Female U50,488 treated RGS7^+/+^ and RGS7^-/-^ mice display no genotypic difference in latency to jump at a 3^oC^ (t _(11)_ =0.1858, p=0.856) **(S. Figure 1H)**. Together our data shows that RGS6 but not RGS7 modulates anti-nociceptive and nocifensive KOR signaling.

### RGS6 nor RGS7 modulates KOR-mediated sedation

We have shown that RGS6 modulates KOR-induced anti-nociceptive behavior, however it is unknown whether this is specific to anti-nociception or if RGS6 or RGS7 can affect other KOR-mediated behaviors. An accelerating rotarod has commonly been used to evaluate motor incoordination or sedative effects induced by opioids, including KOR agonists (Kieffer, 1999) **(Figure 5A)**. First, we assessed if there were any sex differences present in C57BLJ6 mice treated with U50,488 and found similar baseline levels in males and females (t _(11)_ = 0.4818, p=0.639) **(Figure 5B)** and that U50,488 induces significant levels of sedation (time F _(4, 44)_ = 27.26, p<0.0001) at 10 and 30 minutes after injection in both sexes **(Figure 5C).** Male RGS6^+/+^ and RGS6^-/-^ mice showed no difference between genotypes at baseline (t _(14)_ =0.0516, p=0.9596) **(Figure 5D)** nor in sedative recovery (F _(1, 14)_ = 0.1410, p=0.713) **(Figure 5E)**. Similarly, female RGS6^+/+^ and RGS6^-/-^showed no difference at baseline (t _(14)_ =0.3095, p=0.7615) **(Figure 5F)** or sedative recovery (F _(1, 14)_ =0.0001194, p=0.9914) **(Figure 5G)**. Altogether this suggests that both sexes of RGS6^-/-^ mice have no deficits in KOR-mediated sedation and recovery compared to RGS6^+/+^ mice. Testing male RGS7^+/+^ and RGS7^-/-^ mice also demonstrated no differences in their baseline (t _(11)_ =0.293, P=0.775) **(Figure 5H)** nor sedative recovery to a sedative dose of U50,488 (F _(1, 10)_ = 0.3550, p=0.5646) **(Figure 5I)**. Female RGS7 mice were similar to males in that there was no phenotypic difference shown in either their baseline (t _(12)_ =0.3942, p=0.7004) **(Figure 5J)** or in KOR-mediated sedation (F _(1, 12)_ = 0.01608, p=0.9012) **(Figure 5K)**. These results indicate that neither RGS6 nor RGS7 modulates KOR-induced sedative effects.

**Figure 5.**
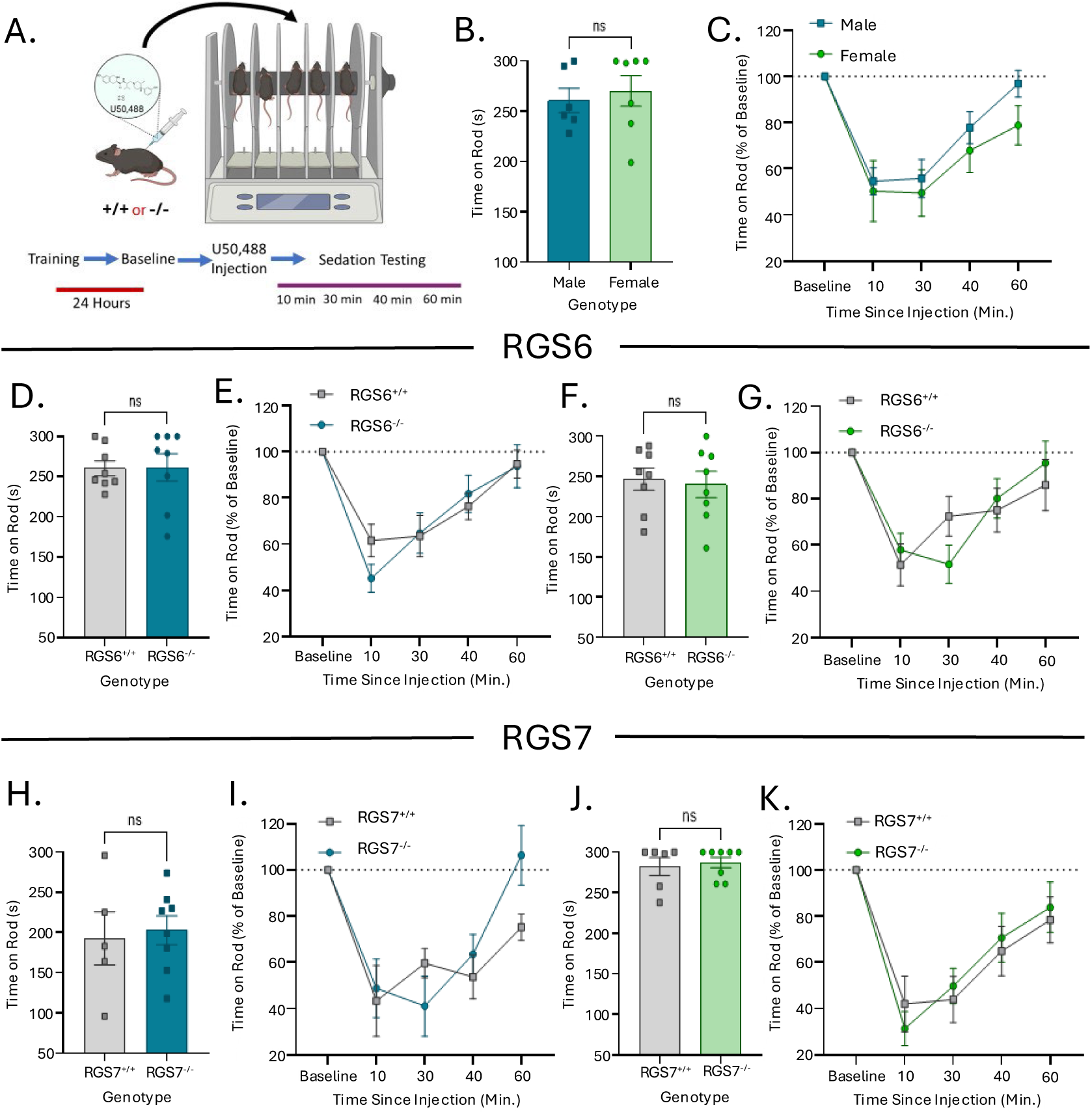
RGS6 and RGS7 on KOR-induced Sedation **A.** Schematic and timeline to test the sedative effect of 15 mg/kg U50,488 on a 3-30 RPM accelerating rotarod in both sexes. Representative colors for each sex are blue for males and green for females. **B.** Average time on the rotarod during baseline for both sexes of C57BLJ6 mice (N=6-7). **C.** Time on the rotarod (% of Baseline) over time after an injection of U50,488 in C57BLJ6 male and female mice. **D.** Average time on the rotarod during baseline for male RGS6^+/+^ and RGS6^-/-^ mice (N=8). **E.** Time on the rotarod (% of Baseline) over time after an injection of U50,488 in male RGS6^+/+^ and RGS6^-/-^ mice. **F.** Average time on the rotarod during baseline for female RGS6^+/+^ and RGS6^-/-^ mice (N=8). **G.** Time on the rotarod (% of Baseline) over time after U50,488 in female RGS6^+/+^ and RGS6^-/-^ mice. **H.** Average time on the rotarod during baseline for male RGS7^+/+^ and RGS7^-/-^ mice (N=5-8). **I.** Time on the rotarod (% of Baseline) over time after U50,488 in male RGS7^+/+^ and RGS7^-/-^ mice. **J.** Average time on the rotarod during baseline for female RGS7^+/+^ and RGS7^-/-^ mice (N=6-8). **K.** Time on the rotarod (% of Baseline) over time after U50,488 in female RGS7^+/+^ and RGS7^-/-^ mice. Data shown as means ± SEM.

### RGS6 and RGS7 do not display compensation in KOR-mediated behaviors in a double knockout model

Both RGS6 and RGS7 are highly co-expressed in Oprk1^+^ cells, therefore we determined if there was compensation between RGS6 and RGS7 using a double knockout (dKO) of RGS6 and RGS7 (RGS6^-/-^ /RGS7^-/-^). Using the cold plate assay, we assessed the collaborative contribution of RGS6 and RGS7 on noxious cold hypersensitivity and nocifensive behaviors in male and female mice. We found that compared to saline-induced cold nocifensive jumping, U50,488-treated male RGS7^-/-^ littermates show significant elevation in nocifensive responses, with no effect on dKO nocifensive jumps (treatment (F _(1, 28)_ = 6.420. p=0.0172; genotype (F _(1, 28)_ = 10.35, p=0.0033) **(Figure 6A).** Examination of latency to noxious cold induced nocifensive jumping revealed that male dKO mice showed a higher latency to respond compared to RGS7^-/-^, confirming the blunting of KOR-induced hypersensitivity in male dKO mice (t _(17)_ =3.568, p=0.0024) **(S. Figure 2A)**. A similar phenotype was also present in female as dKO mice showed blunted nocifensive behavior compared to RGS7^-/-^ mice when exposed to U50,488 at 3^oC^ (treatment F _(1, 29)_ = 13.89, p=0.0006; genotype F _(1, 29)_ = 4.246, p=0.0461) **(Figure 6B)**. Female dKO mice showed no effect of genotype on latency of U50,488-dependent jumps compared to RGS7^-/-^ mice (t _(25)_ =0.025, P=0.981) **(S. Figure 2B)**. To better assess compensation between RGS6 and RGS7 on noxious cold hypersensitivity, we compared the average fold change in U50,488 elicited jumps between RGS6^-/-^/RGS7^-/-^ dKO and global RGS6^-/-^ mice. There were no significant differences between RGS6^-/-^ /RGS7^-/-^ dKO and global RGS6^-/-^ mice of both sexes (genotype F _(1, 36)_ = 0.7406, p=0.3951; sex F _(1, 36)_ = 0.8479, p=0.3633) **(Figure 6C)**. Both global RGS6^-/-^ and RGS6^-/-^/RGS7^-/-^ dKO mice show similar blunting of KOR-dependent hypersensitivity and nocifensive behaviors to noxious cold, implicating this phenotype as being due to lack of RGS6.

**Figure 6.**
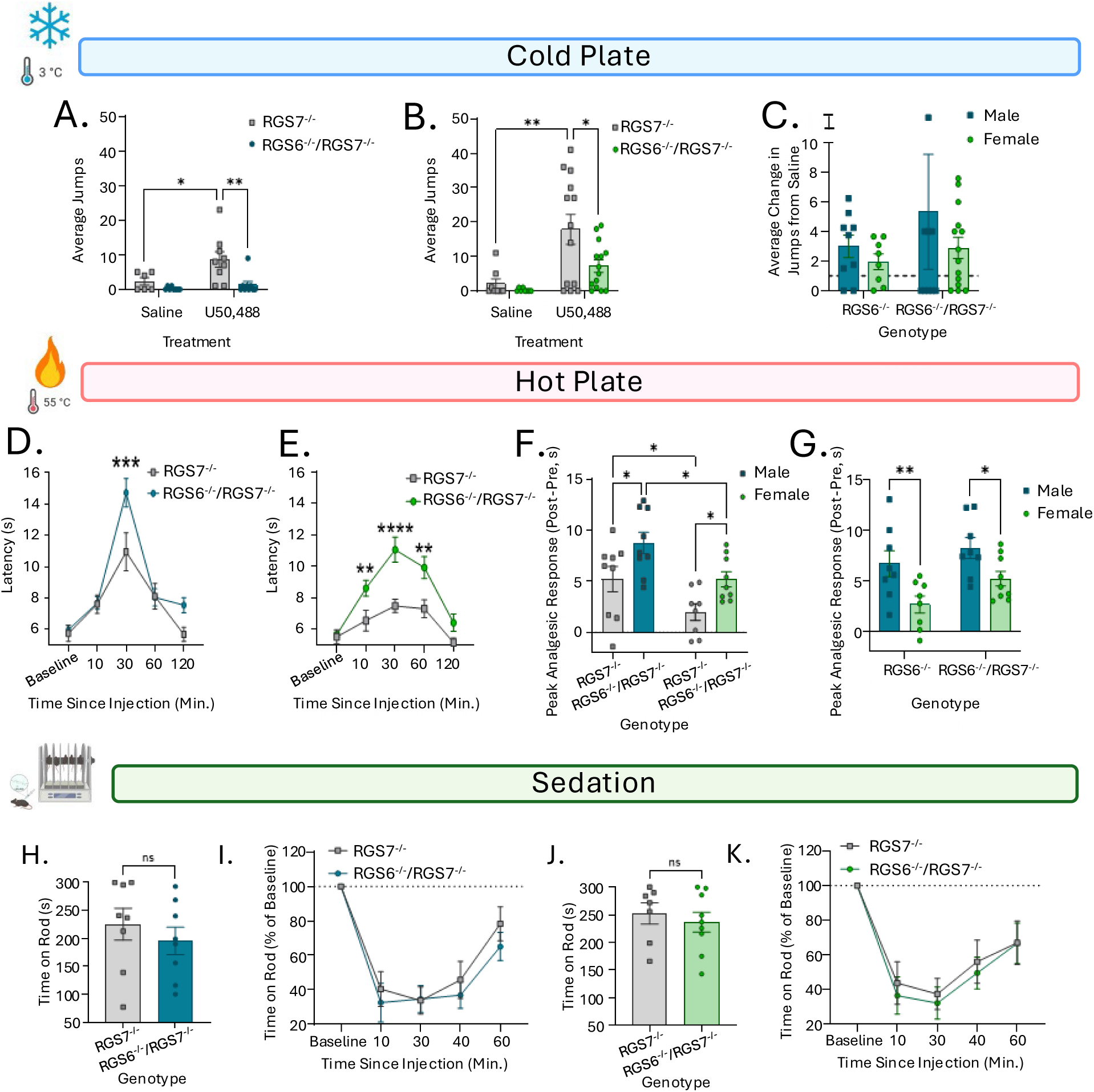
Double knockout (dKO) of RGS6 and RGS7 on KOR-mediated behaviors. Colors assigned to males and females are blue and green respectively. **A.** Average number of jumps after an injection of saline or U50,488 in male RGS6^-/-^/RGS7^-/-^ dKO mice and RGS7^-/-^ littermates (N= 5-9). **B.** Average number of jumps after an injection of saline or U50,488 in female dKO mice and RGS7^-/-^ littermates (N= 13-14). **C.** Comparison of average fold change in noxious cold induced jumps in male and female RGS6^-/-^ and dKO mice. Dotted line at 1 indicates no change. **D.** Average latency to show a pain response in a hot plate assay overtime in male dKO and RGS7^-/-^ littermates (N=9). **E.** Average latency to show a pain response in a hot plate assay over time in female dKO mice and RGS7^-/-^ littermates (N=8-9). **F.** Comparison of peak analgesic response between RGS7^-/-^ and dKO mice in both sexes. **G.** Comparison of peak analgesic response between male and female RGS6^-/-^and dKO mice. **H.** Average time on the rotarod during baseline for male dKO mice and RGS7^-/-^ mice (N=8). **I.** Time on the rotarod (% of Baseline) over time after an injection of U50,488 in male dKO mice and RGS7^-/-^ mice. **J.** Average time on the rotarod during baseline for female dKO and RGS7^-/-^ mice (N=7-9). **K.** Time on the rotarod (% of Baseline) over time after U50,488 treatment in female dKO and RGS7^-/-^ mice. Data shown as means ± SEM; * p≤0.05, ** p≤0.01, *** p≤0.001, **** p<0.0001.

Assessment of KOR-mediated thermal anti-nociception in male RGS6^-/-^/RGS7^-/-^ dKO and RGS7^-/-^ littermates both showed an increase in latency over time, however RGS6^-/-^/RGS7^-/-^ dKO mice displays an elevated anti-nociceptive phenotype compared to RGS7^-/-^ as shown in an increased latency to show a pain response (time F _(4,64)_ = 39.20, p<0.0001; genotype F _(1, 16)_ = 4.669, p=0.0462; interaction F _(4, 64)_ = 3.567, p=0.0109) **(Figure 6D)**. U50,488-treated female RGS6^-/-^

/RGS7^-/-^ dKO mice exhibited a significantly higher latency and an extended time scale of anti-nociception compared to RGS7^-/-^ mice, with RGS7^-/-^ mice showing no change from baseline (genotype F _(1, 14)_ = 34.04, p<0.0001; time F _(4, 56)_ = 18.38, p<0.0001; interaction F _(4, 56)_ = 2.884, p=0.0305) **(Figure 6E)**. Comparison of the peak analgesic response between male and female RGS6^-/-^/RGS7^-/-^ dKO and RGS7^-/-^ counterparts, revealed that RGS6^-/-^/RGS7^-/-^ dKO of both sexes show increased analgesia compared to RGS7^-/-^ but that female mice displayed dampened thermal analgesia regardless of genotype (genotype F _(1, 31)_ = 11.68, p=0.0018; sex F _(1, 31)_ =11.76, p=0.0017) **(Figure 6F)**. To determine if the dKO unmasked any compensatory phenotypic effects, we compared the peak analgesic response in both sexes of RGS6^-/-^and RGS6^-/-^/RGS7^-/-^ dKO mice. It was found that while both RGS6 and dKO showed sex dependent effect of KOR, there was no difference in the peak analgesic response of KOR activation on noxious thermal thermal stimuli in RGS6^-/-^and RGS6^-/-^/RGS7^-/-^ dKO mice (sex F _(1, 29)_ =12.97, p=0.0012; genotype F _(1, 29)_ =4.397, p=0.0448) **(Figure 6G)**. This implicates a lack of compensatory effects in the anti-nociceptive or nocifensive effects of KOR activation. Additionally, we evaluated both sexes of RGS6^-/-^/RGS7^-/-^and RGS7^-/-^ KO mice on KOR sedation. Male RGS6^-/-^/RGS7^-/-^ and RGS7^-/-^ dKO mice displayed no baseline difference (t _(14)_ =0.793, p=0.4408) **(Figure 6H)** and similar sedative recovery levels over time (time F _(4, 56)_ =41.64, p<0.0001; genotype F _(1, 14)_ =0.4420, p=0.5169) **(Figure 6I)**. Female RGS6^-/-^/RGS7^-/-^ and RGS7^-/-^ dKO mice showed equivalent baseline values (t _(14)_ =0.5936, p=0.5623) **(Figure 6J)** and comparable levels of KOR-induced sedation over time (time F _(4, 52)_ = 29.44, p<0.0001; genotype F _(1, 13)_ = 0.1281, p=0.7261) **(Figure 6K)**. Overall, elimination of both RGS6 and RGS7 showed no difference from global RGS6^-/-^ mice on all tested KOR-mediated behaviors suggesting that RGS6 and RGS7 do not compensate for one another.

### Sex dependent regulation of peripheral KOR-mediated sensory processing and anti-nociception by RGS6

Given that RGS6 modulated KOR-induced noxious cold hypersensitivity and nocifensive responses, we investigated if RGS6 modulates this behavioral phenotype centrally or peripherally. We used two highly specific and peripherally restricted KOR agonists, ICI 204,488 and FE 200665 on RGS6^+/+^ and RGS6^-/-^ mice of both sexes in a 3^oC^ cold plate assay **(Figure 7A)**. In male RGS6^+/+^ and RGS6^-/-^ mice treated with ICI 204,488, we found significantly increased nocifensive behavior in both genotypes when treated with ICI 204,488 compared to saline (treatment F _(1, 28)_ = 24.05, p<0.0001; genotype F _(1, 28)_ = 0.3257, p=0.5728) **(Figure 7B)**. ICI 204,488 treated male RGS6^+/+^ and RGS6^-/-^ mice display comparable latency to the nocifensive cold response (t _(12)_ =0.02863, p=0.9776) **(S. Figure 3A).** Female RGS6^+/+^ treated with ICI 204,488 also displayed increased number of jumps on the cold plate assay, whereas RGS6^-/-^ mice showed lower responsivity with fewer jumps recorded (treatment F _(1, 32)_ = 17.79, p=0.0002; genotype F _(1, 32)_ = 9.904, p=0.0036) **(Figure 7C)**. Female latency to show ICI 204,488 induced nocifensive responses to cold showed no significant increase in latency between WT and KO though there is a trend towards increased latency (t _(18)_ =1.772, p=0.0934) **(S. Figure 3B)**. FE 200665, an additional peripherally restricted KOR agonist, was used to test if this phenotype was also present when treated with a peptidergic compound. We found both male RGS6^+/+^ and RGS6^-/-^ mice showed enhanced nocifensive jumping behavior but there was no effect of genotype similar to what we saw with RGS6 males treated with ICI 204,488 (treatment F _(1, 28)_ = 12.03, p=0.0017; genotype F _(1, 28)_ = 0.4543, p=0.5058) **(Figure 7D)**. FE 200665 treated mice exposed to noxious cold showed equivalent latency to jump in both genotypes (t _(11)_ =1.428, p=0.1811) **(S. Figure 3C)**. Female RGS6^+/+^ and RGS6^-/-^ mice treated with FE 200665 on a 3^oC^ cold plate exhibited elevated nocifensive jumping compared to saline, with female RGS6^-/-^ mice displaying dampened responsivity to KOR-dependent noxious cold nocifensive behaviors (treatment F _(1, 30)_ = 30.24, p=<0.0001; genotype F _(1, 30)_ = 6.892, p=0.0135) **(Figure 7E)**. Despite female RGS6^-/-^ mice showing decreased jumping behavior compared to RGS6^+/+^ mice, there was no effect of genotype on latency to the first jump (t _(16)_ =0.3650, p=0.7199) **(S. Figure 3D).** Both ICI 204,488 and FE200665 induced cold hypersensitivity upon KOR activation, indicating that this behavioral output is at least partially driven by peripheral KORs. However, in response to peripherally restricted KOR agonists, the previously observed diminished noxious cold hypersensitivity in RGS6^-/-^ mice treated with central KOR agonists was gone in males but persisted in female RGS6^-/-^ mice.

**Figure 7.**
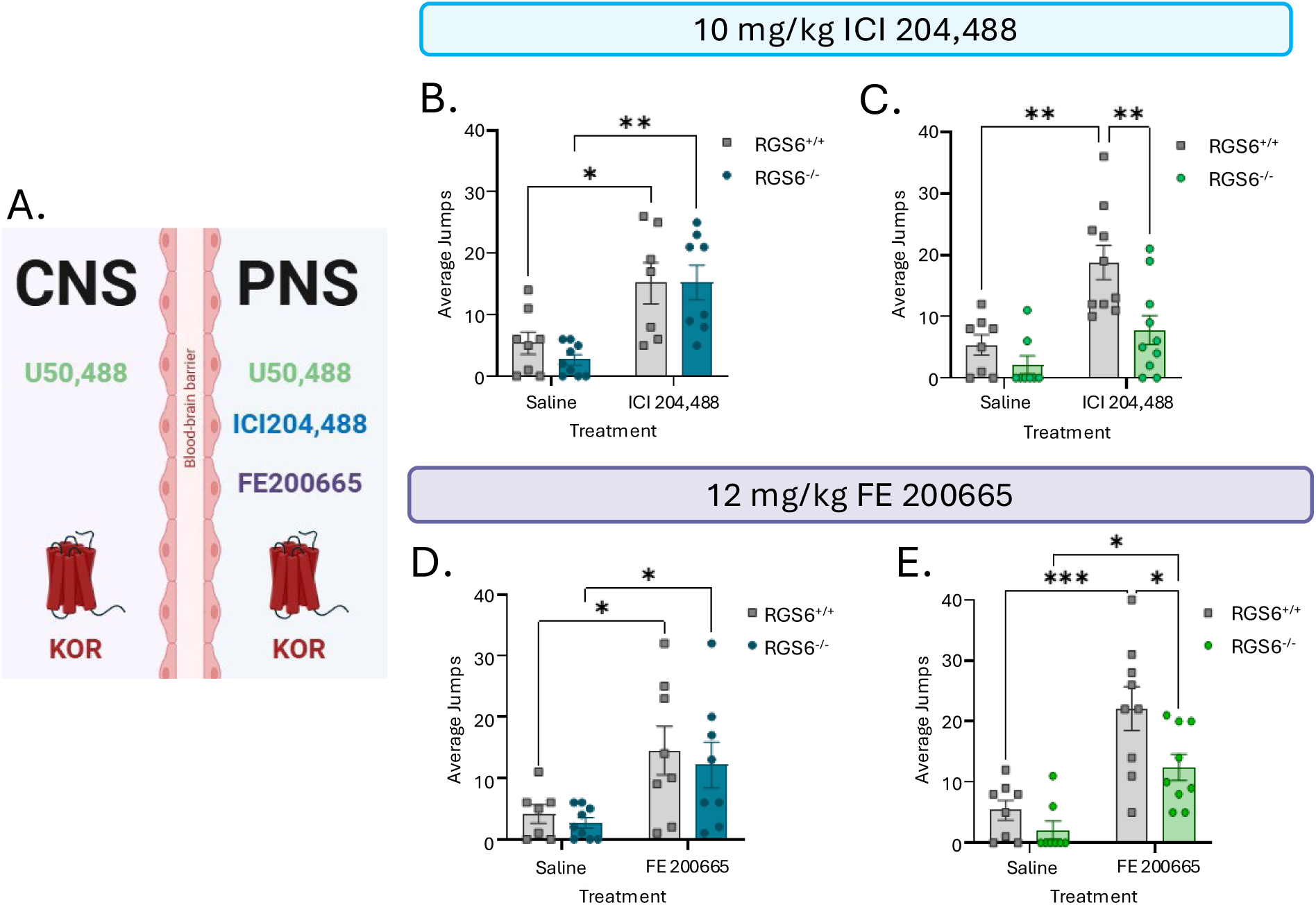
Peripherally restricted KOR agonists on noxious cold hypersensitivity in both sexes of RGS6^+/+^ and RGS6^-/-^ mice. Colors assigned to males and females are blue and green respectively. **A.** Schematic illustrating the KOR agonists used and where their mechanism of action is located. **B.** Average number of KOR-induced nocifensive jumps on a 3^oC^ cold plate after an injection of saline or 10 mg/kg ICI 204,488 in male RGS6^+/+^ and RGS6^-/-^ mice (N=7-8). **C.** Average number of jumps on a 3^oC^ cold plate after an injection of saline or ICI 204,488 in female RGS6^+/+^ and RGS6^-/-^mice (N=10). **D.** Average number of nocifensive jumps on a 3^oC^ cold plate after an injection of saline or 12 mg/kg FE 200665 in male RGS6^+/+^ and RGS6^-/-^ mice (N=8). **E.** Average numbers of jumps on a 3^oC^ cold plate after an injection of saline or FE 200665 in female RGS6^+/+^ and RGS6^-/-^mice (N=9). Data shown as means ± SEM; * p≤0.05, ** p≤0.01.

We have shown that RGS6 modulates KOR-induced anti-nociception in response to U50,488, a central and peripherally circulating agonist, so next we wanted to assess the contribution of RGS6 to mechanical nociception using peripherally restricted KOR agonists in both sexes. Male RGS6^+/+^ and RGS6^-/-^ mice treated with ICI 204,488 showed a significant increase in MWTs at the peak analgesic time point compared to baseline, with no effect of genotype (time F _(5, 75)_ = 11.51, p<0.0001; genotype F _(1, 15)_ = 1.123, p=0.3061; Insert t _(16)_ =0.4753, p=0.6410) **(Figure 8A)**. Female ICI 204,488 treated RGS6^+/+^ and RGS6^-/-^ mice both displayed enhanced MWT, with RGS6^-/-^ mice showing enhanced anti-nociceptive effects to mechanical stimuli (time F _(5, 80)_ = 15.93, p<0.0001; genotype F _(1, 16)_ = 9.570, p=0.007) **(Figure 8B)**. Comparison of MWT at the peak analgesic time points showed that RGS6^-/-^ had higher MWTs on average than RGS6^+/+^ mice (Insert t _(16)_ =2.217, p=0.0415) **(Figure 8B, Insert)**. FE 200665 treated male RGS6^+/+^ and RGS6^-/-^ mice show an increase in their MWTs over time, however there was no difference in thresholds between genotypes including at the peak analgesic time point (time F _(5, 50)_ = 24.43, p<0.0001; genotype F _(1, 10)_ = 0.7751, p=0.3993; Insert t _(10)_ =0.007110, p=0.9945) **(Figure 8C)**. Female RGS6^+/+^ and RGS6^-/-^ mice treated with FE 200665 also showed increased MWT overtime after injection, with RGS6^-/-^ mice displaying heightened MWT compared to RGS6^+/+^ mice (time F _(5, 50)_ = 18.28, p<0.0001; genotype F _(1, 10)_ = 27.44, p=0.0004; interaction F _(5, 50)_ = 2.427, p=0.0479) **(Figure 8D)**. Female RGS6^-/-^ mice showed increased anti-nociceptive effects at the peak analgesic time point (t _(10)_ =3.378, p=0.007) **(Figure 8D, Insert)** Both peripheral KOR agonists induced no phenotypic difference in male RGS6^-/-^ mice compared to RGS6^+/+^, but did induce a substantial increase in KOR mechanical anti-nociception in female RGS6^-/-^ compared to female RGS6^+/+^ mice.

**Figure 8.**
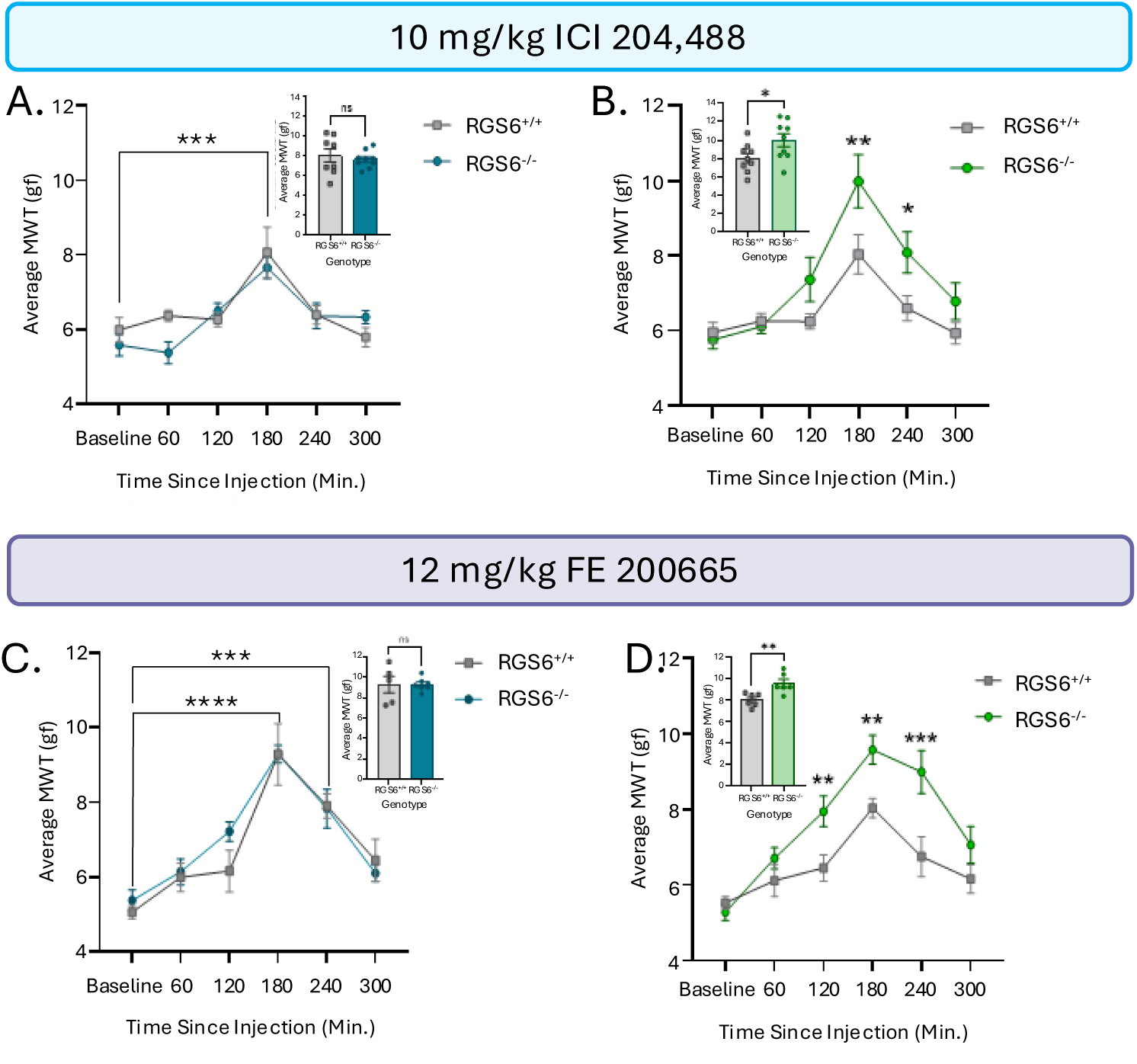
ICI 204,488 and FE 200665-induced Von Frey in RGS6^+/+^ and RGS6^-/-^ mice of both sexes. Colors assigned to males and females are blue and green respectively. **A**. Average MWT after 10 mg/kg ICI 204,488 injection over time in male RGS6^+/+^ and RGS6^-/-^ mice. Insert is the average MWT of male RGS6^+/+^ and RGS6^-/-^ at the peak analgesic response point of 180 minutes after injection (N=8-9). **B.** Average MWT after ICI 204,488 injection over time in female RGS6^+/+^ and RGS6^-/-^ mice. Insert is the average MWT of female RGS6^+/+^ and RGS6^-/-^ at peak response time (N=9). **C.** Average MWT after 12 mg/kg FE 200665 injection over time in male RGS6^+/+^ and RGS6^-/-^ mice. Insert is the average MWT of male RGS6^+/+^ and RGS6^-/-^ at peak response time (N=5-6). **D.** Average MWT after FE 200665 injection over time in female RGS6^+/+^ and RGS6^-/-^ mice. Insert is the average MWT of female RGS6^+/+^ and RGS6^-/-^ at the peak response time (N=6-5). Data shown as means ± SEM; * p≤0.05, ** p≤0.01, *** p≤0.001, **** p<0.0001.

## DISCUSSION

KOR activation produces robust anti-nociception via G protein mediated signaling, but the molecular diversity of intracellular modulators that shape KOR signaling and its behavioral consequences is limited. In this study we demonstrated that RGS6, a main regulator of Gα_i/o_ signaling, modulates multiple modalities of KOR-induced anti-nociception and nocifensive behaviors, with no effect on KOR-dependent sedation. RGS6 knockout mice display thermal and mechanical anti-nociception, enhancing analgesic output and blunting noxious cold hypersensitivity and nocifensive behaviors upon KOR activation. Interestingly, we found that this effect was specific to RGS6 as there is no compensation from RGS7 despite its wide expression patterns in KOR neurons. Furthermore, female RGS6^-/-^ mice exhibited enhanced mechanical anti-nociception and attenuated noxious cold hypersensitivity upon administration of peripherally acting KOR agonists while male mice failed to exhibit this heightened KOR-mediated anti-nociceptive sensitivity. These findings present new evidence demonstrating that RGS6 acts as a negative regulator of KOR signaling that regulates anti-nociception and may differentially modulate KOR in central and peripheral nociceptive circuits.

RGS6 and RGS7 are highly expressed in several areas essential for pain perception, both in spinal and DRG nociceptive circuits. Here we show that not only is RGS6 and RGS7 highly enriched in KOR expressing neurons within these circuits, but that lack of RGS6 enhances KOR-induced anti-nociception to noxious thermal and mechanical stimuli. Even though these RGS proteins are expressed in the same cells and accelerate GTP hydrolysis on Gα_i/o_, they appear to have specificity towards KOR-G protein signaling. In other GPCR systems, RGS6 and RGS7 have also been shown to display distinct receptor-dependent signaling (Luo et al., 2024; Masuho et al., 2020). For example, in hippocampal neuronal cultures RGS7^-/-^ mice display enhanced the deactivation of GABA_B_R-G protein-gated inwardly rectifying potassium (GIRK) signaling, whereas RGS6^-/-^ neurons show no impact GABA_B_R-GIRK signaling (Ostrovskaya et al., 2014). Furthermore, RGS7 ablation slows activation of GIRK signaling in response to 5HT_1A_R, whereas loss of RGS6 prolongs this activation of 5HT_1A_R-GIRK response (Luo et al., 2024). In part this may be due to the distinct sets of binding partners that establish differential complex formation contributing to their divergent signaling roles. For example, both can associate with core partners such as Gβ5, but they show different preferences and dependencies for accessory proteins like R7BP and GPR158/GPR179, which affect their subcellular localization and membrane targeting (Drenan et al., 2005; Hu and Wensel, 2002; Martemyanov et al., 2005; Orlandi et al., 2012; Sandiford and Slepak, 2009; Song et al., 2006). This formation positions RGS proteins in partially non-overlapping signaling microdomains that help shape the targeting of distinct GPCR-G protein signaling pathways. Our results show that even within the R7 RGS family, RGS proteins differentially regulate KOR-Gα_i/o_ and may have implications on the development of novel therapeutic analgesic treatments.

KOR agonists are well known to induce significant levels of sedation in both preclinical mice and human studies, which has limited its therapeutic potential. Here we found that neither single nor double KO of RGS6 and RGS7 altered KOR-induced sedation, indicating that these RGS proteins are involved in this behavioral effect. Instead, KOR-RGS6 modulation is selective to specific KOR-mediated behavioral outcomes as RGS6 modulates KOR anti-nociception but not sedation. This behavioral response is likely due to differences in signaling pathways. Canonically, KOR-dependent sedation has long been attributed to β-Arrestin signaling. Evidence from studies investigating the development of G protein biased agonists have shown that β-Arrestin is necessary for KOR-induced sedation (Brust et al., 2016; Dunn et al., 2019; Endoh et al., 1999). Knockout studies of β-Arrestin2 have exhibited enhanced motor coordination and sedative recovery compared to wildtype mice when treated with KOR agonists, demonstrating that β-Arrestin plays an essential role in mediating KOR-induced sedation (White et al., 2015). Together, these findings support a model in which RGS6 regulates KOR-mediated G protein signaling, with minimal impact on β-arrestin pathways, thereby favoring anti-nociceptive actions over sedative side effects.

We have shown that lack of RGS6 modulates KOR-induced anti-nociception and nocifensive responses to multiple types of noxious stimuli, suggesting that RGS6 may regulate KOR signaling in diverse cell populations rather than being restricted to a single nociceptive cell type. Recent work has suggested that central and peripheral KOR can modulate different types of pain modalities. KOR attenuates mechanical hypersensitivity centrally in spinal circuits and supraspinal areas, and peripherally at the DRGs, peripheral nerves, and epidermis (Ma et al., 2023; Nguyen et al., 2022; Pando et al., 2025; Snyder et al., 2018). Modulation of KOR-induced thermal anti-nociception is primarily modulated at the supraspinal and spinally level depending on the assay used (Craft and Bernal, 2001; Vanderah et al., 2008). Because RGS6^-/-^ mice exhibited altered anti-nociceptive and nocifensive behaviors, our findings implicate that RGS6 may modulate KOR signaling at multiple sites. These primary afferent populations carry distinct molecular markers that encode specific forms of noxious information and have been linked to particular nociceptive modalities. Thus, although we demonstrate that RGS6 is broadly expressed at central and peripheral sites and modulates KOR within KOR⁺ nociceptive circuits, the precise neuronal or primary afferent subtypes through which RGS6 exerts these effects remain to be identified.

KOR has long been implicated to modulate cold somatosensation and somewhat controversially, thermoregulation (Madasu et al., 2021; Rawls et al., 2005). Cold hypersensitivity is frequently associated with neuropathic pain, peripheral nerve injury, and chemotherapy-induced peripheral neuropathy (Cobos et al., 2018; Paton et al., 2022). Its comorbidity with nociceptive pain has heightened interest in further exploring noxious cold sensation as one of the major somatosensations. Cold sensation has been attributed to transient receptor potential (TRP) channels, specifically TRPA1 (TRP Ankyrin 1) and TRPM8 (TRP Melastatin 8), both of which are activated by noxious temperatures below 10^oC^ (Pogorzala et al., 2013). Loss of TRPA1 reduces sensitivity to cold and noxious mechanical stimuli, and this diminished cold sensitivity in TRPA1^⁻/⁻^ mice has been linked to increased constitutive KOR activity in TRPA1-expressing sensory neurons, as pharmacological KOR blockade restores nociceptive responses (Semizoglou et al., 2022). Consistent with this, optogenetic activation of afferent KOR induces nocifensive responses including rapid withdrawal of limbs and excessive jumping, a canonical nocifensive response to noxious stimuli (Snyder et al., 2018). Recent work by Madasu *et al*. (2024) further showed that KOR activation can drive noxious cold hypersensitivity and nocifensive behavior through modulation of TRPA1 channels. In our transgenic models, deletion of RGS7 had no effect, whereas RGS6 deletion significantly blunted KOR-dependent cold hypersensitivity and nocifensive behavior to U50,488, a centrally acting KOR agonist, implicating RGS6 as a key modulator of KOR signaling in noxious cold hypersensitivity. RGS6 expression has recently been linked to decrease with the onset of cold allodynia, sensitivity to non-painful stimuli. In a mouse model investigating transcript regulation in the DRGs after spared nerve injury (SNI), RGS6 was found to be downregulated during the onset of neuropathic cold allodynia (Cobos et al., 2018). This suggests in concert with our results, that RGS6 may modulate cold nociceptive signaling. It is possible that an interaction between TRPA1, KOR, and RGS6 may modulate cold hypersensitivity and nocifensive behaviors, although the precise mechanisms by which TRPA1 controls KOR constitutive activity and how RGS6 influences this process remain to be determined. Several studies have demonstrated that analgesic effects of KOR activation are blunted in females compared to male in both human and rodent studies (Craft and Bernal, 2001; Mogil et al., 2003; Stoffel et al., 2005). This sex difference is due to sequestering of the Gβγ subunits by GRK2 due to phosphorylation by elevated estradiol, thus causing a reduced analgesic effect in females (Abraham et al., 2018). RGS6 was shown to modulate KOR-induced anti-nociception to thermal stimuli, while still maintaining this sex difference when treated with the KOR agonist U50,488 a drug that is both central and peripherally acting. RGS6 acts as a GAP for Gαi/o, resulting in the reassociation of Gα and Gβγ subunits, ultimately fine tuning GPCR responses. Thus, in our RGS6^-/-^ mice there is an increase in availability of free Gβγ which could enhance KOR-mediated anti-nociception despite preserved estradiol–GRK2–dependent sex differences in signaling efficiency.

Through the use of two peripherally restricted KOR agonists, we found that RGS6 modulates KOR-dependent anti-nociception peripherally in females but not male mice. This suggests that KOR-RGS6 signaling displays peripheral nervous system (PNS) specific sex differences. Sex differences in the KOR system have been postulated to be mediated spinally rather than supraspinal, but have not been fully explored in the DRGs of the PNS in response to acute pain (Craft and Bernal, 2001). Profound sex differences in the DRGs have been demonstrated in human and mouse studies at the genetic, mechanistic, and behavioral levels. For example, it has been shown that male and females display differentially expressed genes associated with neuropathic pain in the DRGs (Ray et al., 2022). Furthermore, a study investigating sexually dimorphic pain responses in the DRG found a novel female specific mechanism of hyperalgesia via female selective prolactin signaling (Patil et al., 2019). As RGS6 modulates KOR-induced anti-nociception in the PNS in a female specific manner, it is likely that KOR-RGS6 modulates nociceptive circuits in sexually dimorphic populations of the DRGs. Our data demonstrates that further research is necessary to elucidate KOR-RGS6 sex differences in the periphery.

### Conclusions

Taken together, we show that RGS6 modulates KOR-induced anti-nociception and nocifensive behavior to multiple sensory outputs with no effect on KOR-induced sedation. We found that the R7 RGS family may display receptor specificity and do not display functional redundancy at the KOR. Finally, RGS6 may differentially modulate KOR-induced anti-nociception and nocifensive responses on primary afferent population differently in central versus peripheral nociceptive circuits. Further research is necessary to elucidate the expression patterns of RGS6 and KOR in nociceptive populations and the mechanisms mediating RGS6 specific modulation of KOR-induced behaviors. These findings provide essential information on the intracellular modulators that shape KOR signaling and may aid in the development of novel analgesic drugs and therapeutics.

## ACKNOWLEDGEMENTS

We thank Dr. Kirill A. Martemyanov and Dr. Kevin Wickman for providing the RGS7 and RGS6 transgenic lines, respectively. We thank Sara Gavagan and Leen Jawhar for their technical assistance with cold plate and sedation behavioral assays. We also thank Dr. Tagide deCarvalho for her assistance and use of the College of Natural and Mathematical Sciences imagining core.

## Blount et al. 2026. Supplemental Figure 1-3

**Supplemental Figure 1.**
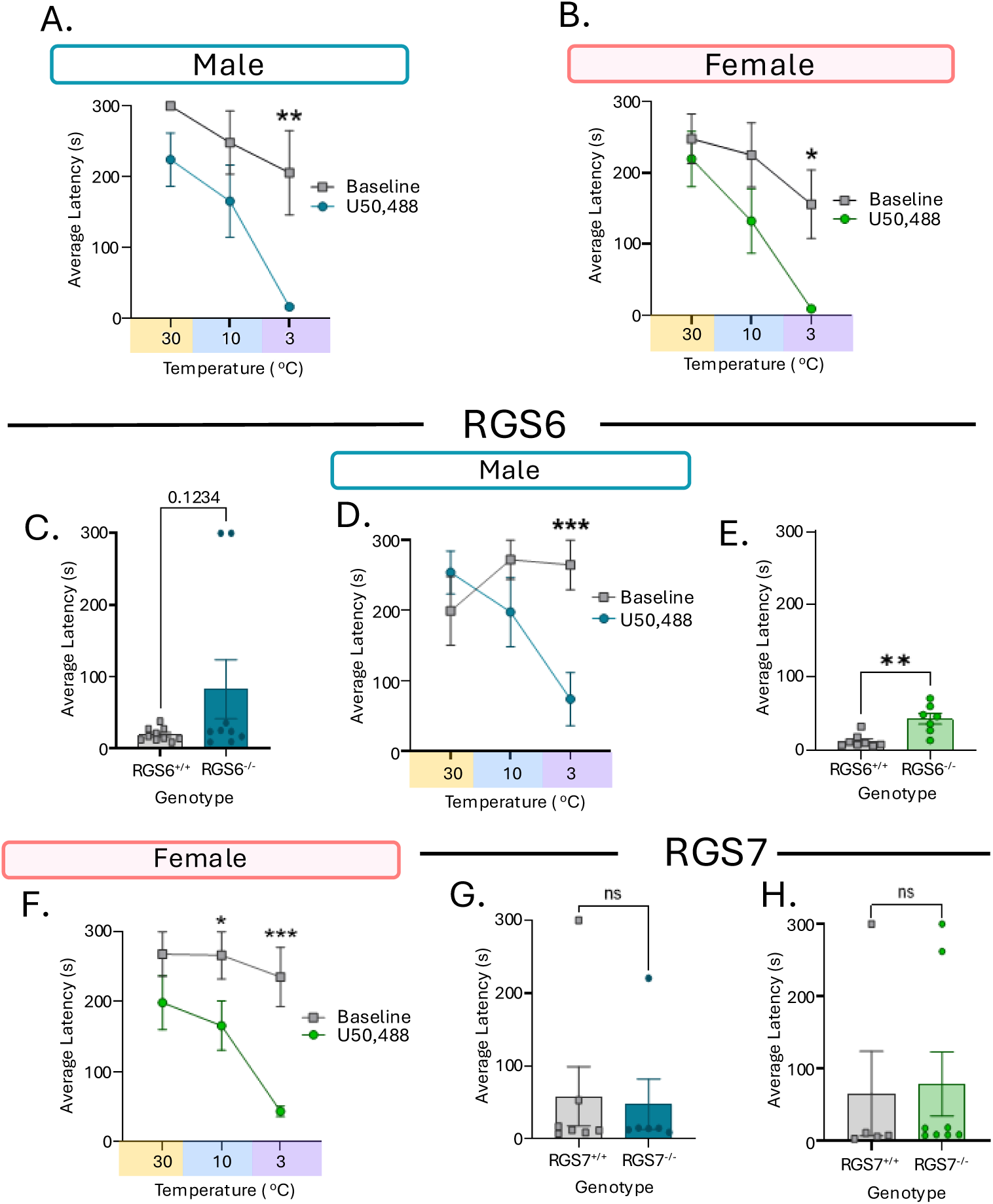
RGS6 and RGS7 on the latency to show KOR-induced noxious cold hypersensitivity and nocifensive behavior. Colors assigned to males and females are blue and green respectively. **A**. Average latency (seconds) to jump in male C57BLJ6 mice when treated with 5 mg/kg U50,488 compared to baseline on a temperature gradient of 30^oC^, 10^oC^, & 3^oC^(N=6-9). **B.** Average latency (seconds) to jump in female C57BLJ6 mice when treated with U50,488 compared to baseline on a temperature gradient (N=8). **C.** Average latency to jump in male RGS6^+/+^ and RGS6^-/-^ mice (N=10-9). **D.** Average latency to jump in male RGS6^+/+^ and RGS6^-/-^mice when treated with U50,488 compared to baseline on a temperature gradient. **E.** Average latency to jump in female RGS6^+/+^ and RGS6^-/-^ mice (N=8-7). **F.** Average latency to jump in female RGS6^+/+^ and RGS6^-/-^ mice when treated with U50,488 compared to baseline on a temperature gradient. **G.** Average latency to jump in male RGS7^+/+^ and RGS7^-/-^ mice (N=7-6). **H.** Average latency to jump in female RGS7^+/+^ and RGS7^-/-^ mice (N=5-8). Data shown as means ± SEM; * p≤0.05, ** p≤0.01, *** p≤0.001, **** p<0.0001.

**Supplemental Figure 2.**
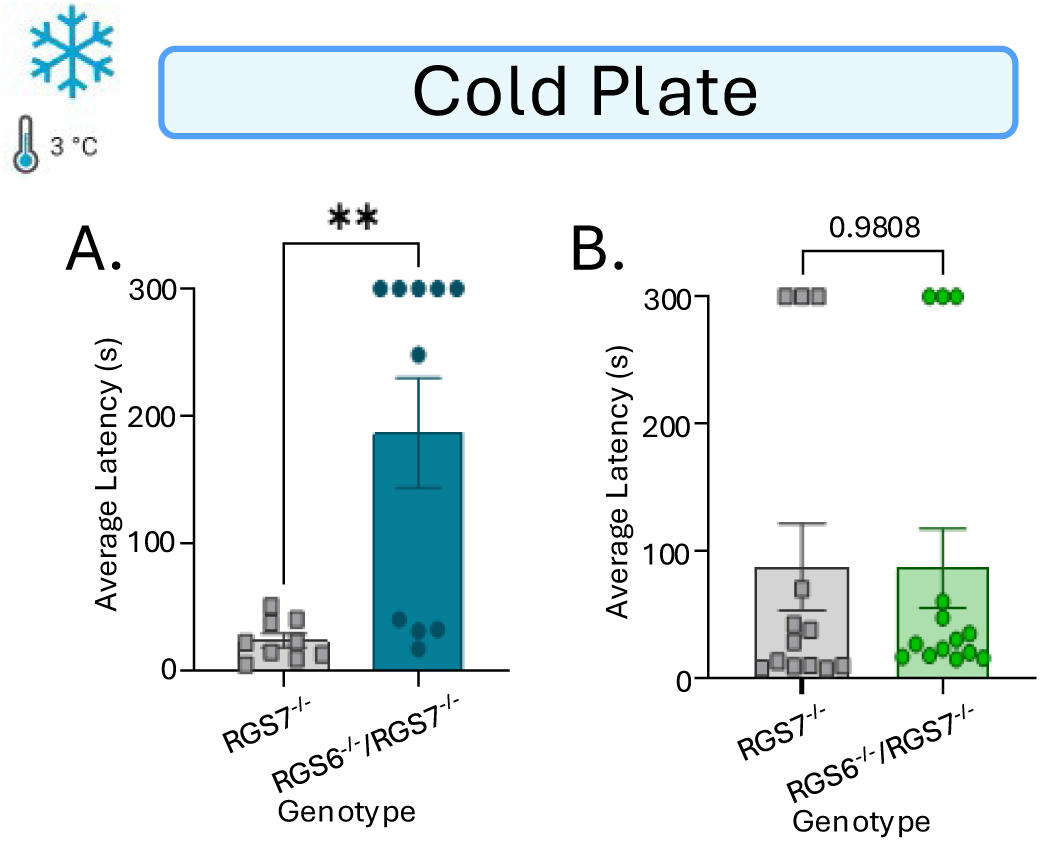
Double knockout of RGS6 and RGS7 on KOR-induced noxious cold hypersensitivity latency. Colors assigned to males and females are blue and green respectively. **A**. Average latency to jump in male RGS6^-/-^RGS7^-/-^ dKOs and RGS7^-/-^ littermates (N=9-10). **B.** Average latency to jump in female RGS6^-/-^RGS7^-/-^dKOs and RGS7^-/-^ littermates (N=13-14). Data shown as means ± SEM; * p≤0.05, ** p≤0.01, *** p≤0.001, **** p<0.0001.

**Supplemental Figure 3.**
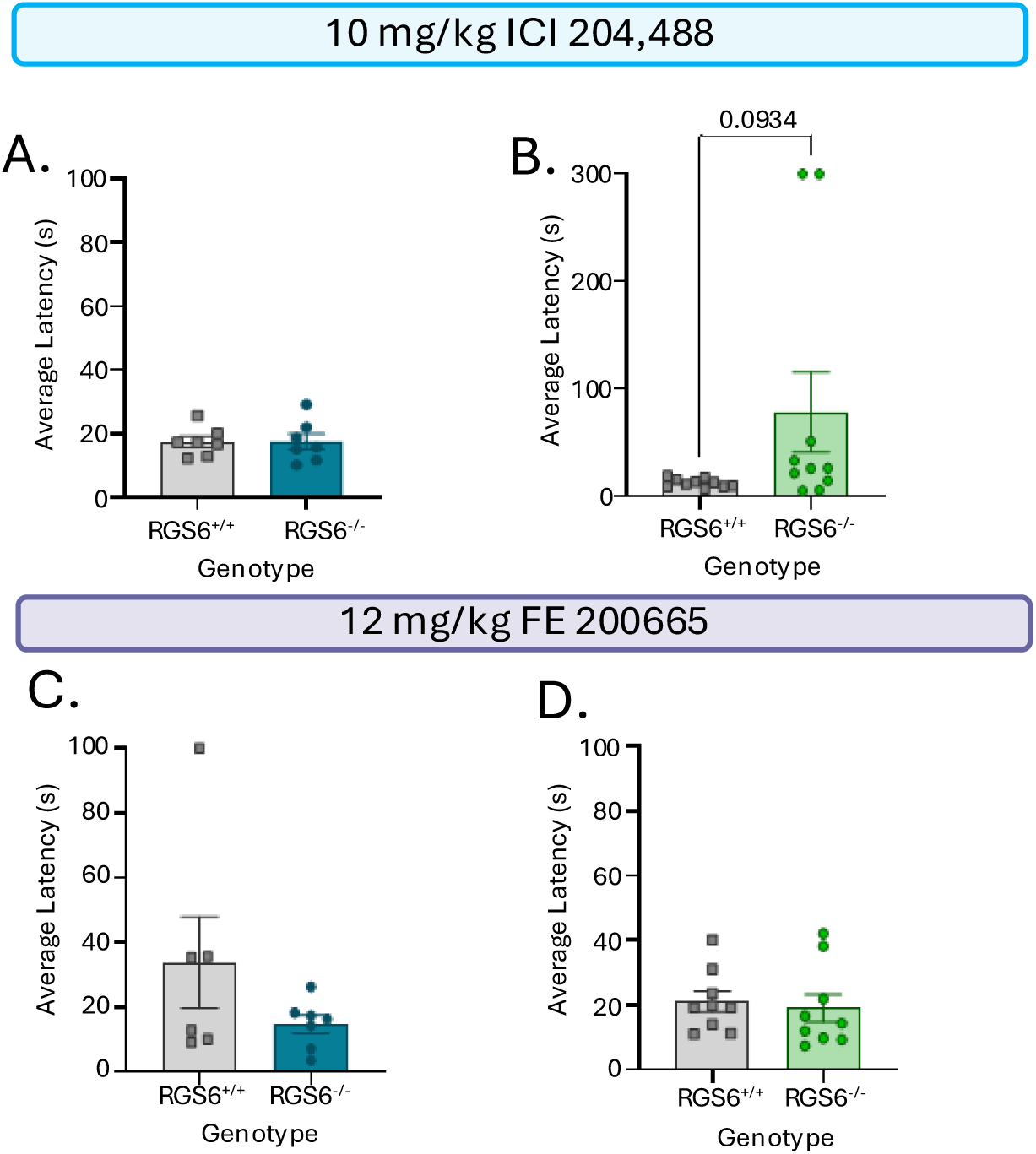
Average latency to jump on the cold plate in RGS6 mice treated with ICI 204,488 or FE 200665**. A.** Average latency to jump in male RGS6^+/+^ and RGS6^-/-^ mice when treated with ICI 204,488 (N=7-8). **B.** Average latency to jump in female RGS6^+/+^ and RGS6^-/-^ mice when treated with ICI 204,488 (N=10). **C.** Average latency to jump in male RGS6^+/+^ and RGS6^-/-^ mice when treated with FE 200665 (N=6-7). **D.** Average latency to jump in female RGS6^+/+^ and RGS6^-/-^mice when treated with FE 200665 (N=9). Data shown as means ± SEM.

